# Experimental Design and Power Calculation in Omics Circadian Rhythmicity Detection

**DOI:** 10.1101/2022.01.19.476930

**Authors:** Wei Zong, Marianne L. Seney, Kyle D. Ketchesin, Michael T. Gorczyca, Andrew C. Liu, Karyn A. Esser, George C. Tseng, Colleen A. McClung, Zhiguang Huo

## Abstract

Circadian clocks are 24-hour endogenous oscillators in physiological and behavioral processes. Though recent transcriptomic studies have been successful in revealing the circadian rhythmicity in gene expression, the power calculation for omics circadian analysis have not been fully explored. In this paper, we develop a statistical method, namely CircaPower, to perform power calculation for circadian pattern detection. Our theoretical framework is determined by three key factors in circadian gene detection: sample size, intrinsic effect size and sampling design. Via simulations, we systematically investigate the impact of these key factors on circadian power calculation. We not only demonstrate that CircaPower is fast and accurate, but also show its underlying cosinor model is robust against variety of violations of model assumptions. In real applications, we demonstrate the performance of CircaPower using mouse pan-tissue data and human post-mortem brain data, and illustrate how to perform circadian power calculation using mouse skeleton muscle RNA-Seq pilot as case study. Our method CircaPower has been implemented in an R package, which is made publicly available on GitHub (https://github.com/circaPower/circaPower).

## Introduction

Circadian rhythms are endogenous ~24 hour oscillations of behavior, physiology, and homeostasis in adaption to the diurnal cycle caused by the earth’s daily rotation. The circadian clock is found in virtually all cells throughout the body and controls oscillations in a wide variety of physiological processes, including sleep-wake cycles, body temperature, and melatonin secretion [1, 2, 3, 4]. From the literature, the mechanism that drives circadian rhythms is a transcriptionial-translational feedback loop encoded by a set of core clock genes [5], including *CLOCK, BMAL1* as the transcriptional activators; and period family (*PER1, PER2, PER3*) and cryptochrome family (*CRY1, CRY2*) as the major inhibitors. In addition to core clock genes, genome-wide transcriptomic studies have revealed additional circadian genes in post-mortem brain [6, 7], skeletal muscle [8], liver [9], and blood [10]. Transcriptomic circadian analyses in human [11], mouse [12], and baboon [13] have shown that the circadian pattern in gene expression could be tissue-specific. Beyond transcriptomic data, circadian rhythmicity was also discovered in other types of omics data including DNA methylation [14], ChIP-Seq (chromatin immunoprecipitation assays with sequencing) [15], proteomics [16], and metabolomics [17]. From epidemiology and animal studies, the disruption in clock and circadian gene expression was found to be linked to diseases including type 2 diabetes [18], cancer [19, 20], sleep [10], major depression disorder [21], aging [6], schizophrenia [7], and Alzheimer’s disease [22].

As the circadian omics studies have become increasingly popular over the years (Figure 1), the experimental design of such circadian omics studies has come into focus [23, 24], where the design refers to the distribution of the collected Zeitgeber time (ZT; standardized diurnal time with ZT0/ZT24 for the beginning of day and ZT12 for the beginning of night). In this paper, we consider two types of sampling design: passive and active sampling design. In passive design, investigators have no control of the collected ZT. Such a passive design is commonly seen in studies with human tissues that are difficult to obtain (e.g., post-mortem brain tissues [6, 21, 7]) and the irregular sampling distribution should be considered in power calculation. In contrast, investigators have full control of the sample collection time in an active sampling design. Such an active design is commonly seen in animal studies [12] or human blood studies [10]. In the literature, 6 time points (every 4 hours) per cycle across one or multiple full cycles have been widely adopted in many actively designed studies [25, 26, 27]. Hughes et al. [28] recommended evenly sampling at least 12 time points per cycle (i.e., every 2 hours) across 2 full cycles. For the ease of discussion, we refer to this type of design as the “evenly-spaced sampling design”. Though these empirical practices and guidelines were presented and well-received, limited quantitative benchmarks are available. To address this, Ness-Cohn et al. [29] developed a user-friendly website, TimeTrial, which allows researchers to explore the effects of experimental design on cycling detection. Although multiple circadian detection methods are allowed, the results are benchmarked through simulation using classification error rate and area under the curve of a ROC curve. However, a statistical method that enables exact power calculation is still lacking. Previous studies have reported the lack of overlapping circadian genes because of smaller number of samples [30, 9], indicating statistical power, i.e., the probability of successfully detecting the underlying circadian pattern, is not fully considered/justified. Thus, an analytical method that allows exact power calculation under different experimental designs is urgently needed. In addition, all these prior works on circadian study design were discussed within the scope of active design, where investigators have control of sample collection time. In the scope for passive design, there are no guidelines in the literature. It is unclear whether and how the irregular ZT distribution will impact the circadian power calculation.

**Fig. 1.**
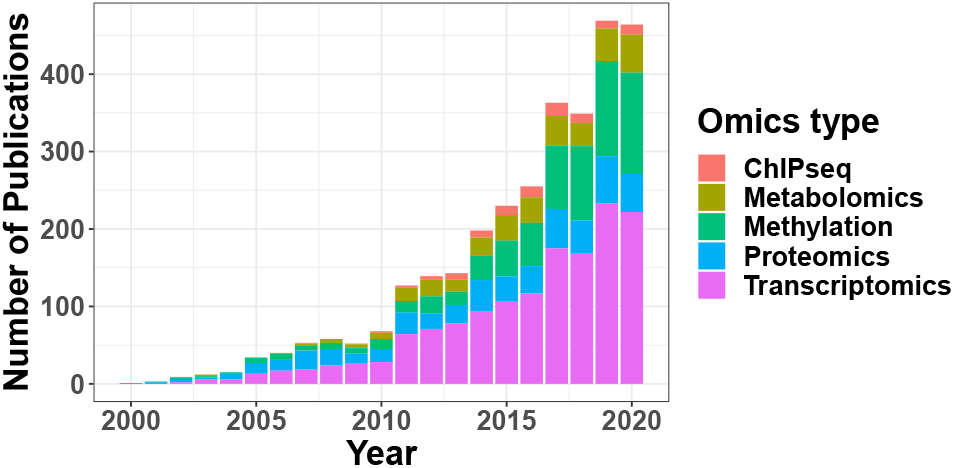
Annual number of publications on PubMed that contain the keywords “circadian/clock” and one of the following omics type: “ChIPseq”, “Metabolomics”, “Methylation”, “Proteomics”, “Transcriptomics”.

To fill i n t hese r esearch g aps, w e p ropose a modelbased approach to accurately calculate the circadian power (namely CircaPower), based on the cosinor model [31, 32]. It assumes the expression level of a gene is a sinusoidal function of the circadian time [31]. Though straightforward, the model is biologically interpretable, and enjoys accurate statistical inferences [32]. In the literature, there are several other algorithms developed for circadian rhythm detection. Lomb-Scargle periodograms [33] and COSOPT [34] (arithmetic linear-regression detrend of time series) are more complicated parametric models. By assuming mixture of multiple cosinor curves with distinct periods, these methods facilitate the detection of oscillating transcripts with irregular shape. ARSER [35] (autoregressive spectral estimation of rhythmicity), RAIN [36] (rhythmicity analysis incorporating nonparametric methods) and JTK CYCLE [37] (nonparametric Jonckheere-Terpstra test) are non-parametric methods, which are free of modeling assumption and more powerful to capture irregular curve shapes. Both parametric and non-parametric algorithms were widely applied in transcriptomic studies, and comparisons of these algorithms have been conducted in several review studies [28, 38, 39, 24]. Although the complex methods have advantages to detect irregular curves beyond cosinor models, the power calculation using these methods are not always feasible since the effect s ize a nd d ata v ariability are not explicitly defined i n t hese m odels. I n a ddition, concerns about the accuracy of the statistical inference for these complex methods have been raised [39]. To be specific, t he p-values generated by many of these methods may not be correct (i.e., do not follow a uniform distribution *UNIF*(0, 1) under the null), implying a potential inflated o r d eflated ty pe I error rate. Therefore, we propose the power calculation framework based on the cosinor model, because of its simplicity and accurate statistical inference. Moreover, the biological rationale for using a cosinor model is that the circadian rhythm is amenable to adapt the cycles in the environment [5], including the day-light cycle, the tides, the phases of the moon, the seasons, etc [40]. Since the day-light cycle is the leading environmental factor that governs circadian rhythms, a cosinor wave model is widely used to mimic the cosinor cycle of the day-light intensity and many previous literatures [6, 7, 41] have used this model to identify biological meaningful findings. We acknowledge that other complex parameters models and nonparametric approaches are also popular with their own unique merit, and exploring circadian power calculation using these complex methods is one of our future directions.

To the best of our knowledge, this is the first theoretical methodology developed for circadian power calculation in omics data. The unique contribution of this paper includes: (i) identifying factors related to the statistical power of circadian rhythmicity detection, including sample size, intrinsic effect size and sampling design; (ii) developing CircaPower, an analytical solution based on a closed-form formula, for fast and accurate circadian power calculation; (iii) demonstrating via simulations that the evenly-spaced sampling design is superior because of its phase-invariant property, which is also corroborated by theoretical proofs; (iv) illustrating how to calculate statistical power and to design a circadian experiment with pilot data via a case study; (v) collecting, calculating, and summarizing the intrinsic effect sizes of existing human a nd a nimal studies, which serves as a useful reference resource when no pilot data is available; and (vi) providing an open-source R package.

The superior performance of our method is demonstrated in comprehensive simulation studies, as well as multiple transcriptomic applications in human and mouse. We demonstrate the performance of CircaPower using continuous gene expression data throughout this manuscript, but our method is also applicable in single biomarker data or other types of continuous omics data, including but not restricted to ChIP-Seq, DNA methylation, proteomics, and metabolomics.

## Methods

The CircaPower framework assumes the relationship of the expression level (continuous data type) of a gene and the Zeitgeber time (ZT) fits a sinusoidal wave curve, and is based on the *F* statistics of a cosinor model [31]. Below we introduce the model notations, the construction of the *F* statistics, the null and alternative distribution of the *F* statistics, the closed-form formula for circadian power calculation, and factors affecting the power calculation of circadian rhythmicity detection.

### Notations and basic model

As illustrated in Figure 2a, denote *y* as the expression value for a gene; *t* as the ZT; *M* as the MESOR (Midline Estimating Statistic Of Rhythm, a rhythm-adjusted mean); *A* as the amplitude. *ω* is the frequency of the sinusoidal wave, where 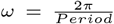. Without loss of generality, we set *period* = 24 hours to mimic the diurnal period. *ϕ* is the phase shift of the sinusoidal wave curve. Whenever there is no ambiguity, we will omit the unit “hours” in period, phase, and other related quantities. Due to the periodicity of a sinusoidal wave, (*ϕ*_1_, *ϕ*_2_) are not identifiable when *ϕ*_1_ = *ϕ*_2_ + 24. Therefore, we will restrict *ϕ* ∈ [0, 24). *ϕ* is not intuitive to read from a sinusoidal wave (see Figure 2a). A closely related quantity is the peak time *t_P_*. The connection between *ϕ* and *t_P_* is that *ϕ* + *t_P_* = 6 ± 24*N*, where *N* is an arbitrary natural number.

**Fig. 2.**
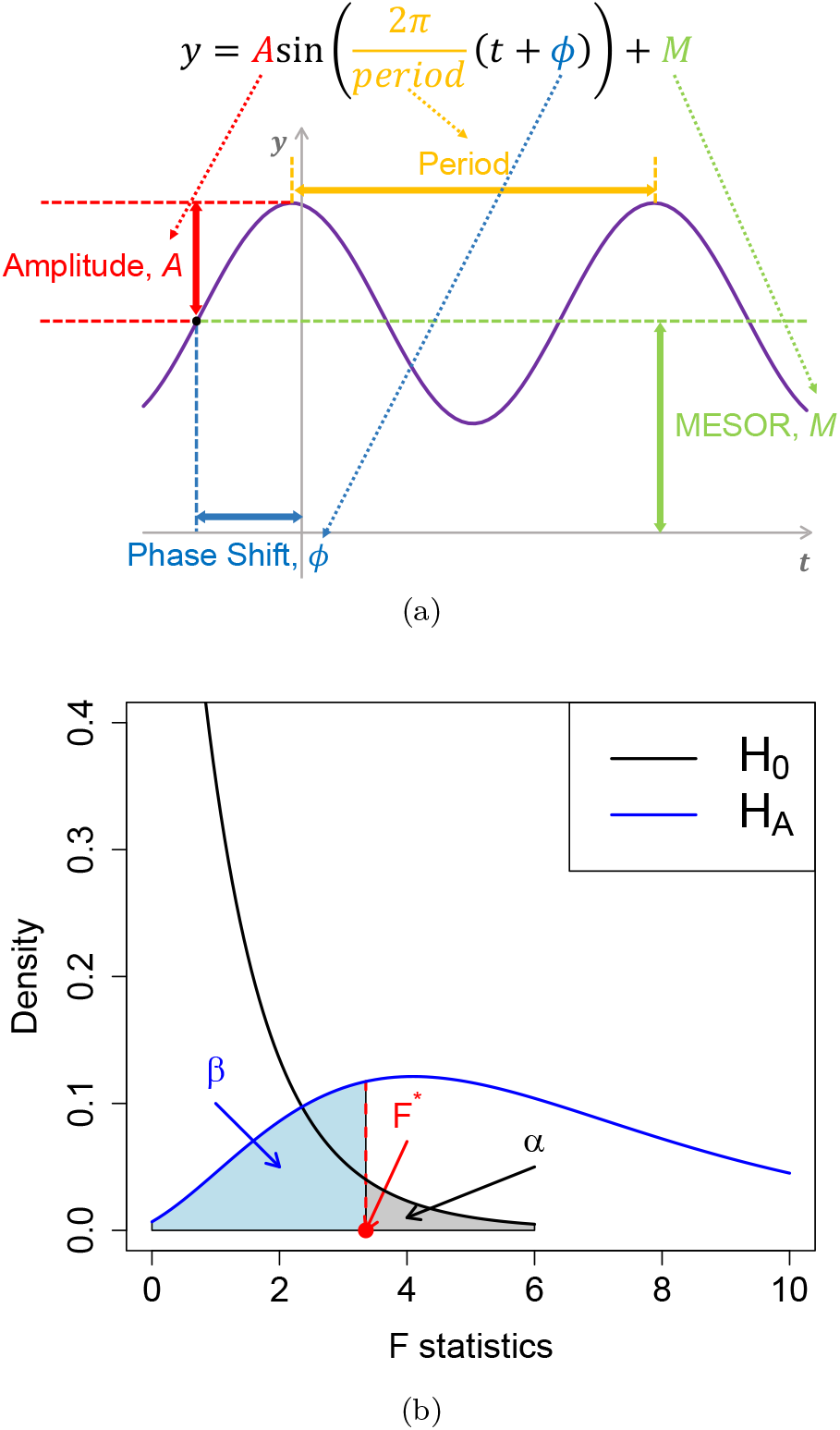
Basic sinusoidal model and the power/type I error discriminatory curve. (a) shows a sinusoidal wave curve underlying circadian rhythmicity power calculation framework. (b) shows the relationship between power and type I control in detecting circadian rhythmicity. The black curve represents the density function of the *F* statistics under the null distribution (no circadian pattern); the blue curve represents the density function of the *F* statistics under the alternative distribution. The red dashed line represents the decision boundary (i.e., *F*\*) such that the type I error rate is controlled at *α* (shaded gray). The corresponding type II error *β* is the area with lightblue color and the detection power is 1 – *β*.

For a given sample *i* (1 ≤ *i* ≤ *n, n* is the total number of samples), denote by *y_i_* the expression value of a gene and *t_i_* the observed ZT. We assume the following sinusoidal wave function:

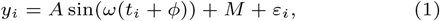

where *ε_i_* is the error term for sample *i*; we assume *ε_i_*’s are identically and independently distributed (*i.i.d*.) from *ε_i_* ~ *N*(0, *σ*^2^), where *σ* is the noise level. To benchmark the goodness of sinusoidal wave fitting, we define the coefficient of determination 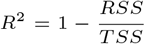, where 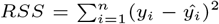, 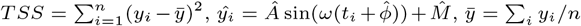, with 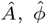, and 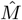 being the fitted value for *A*, *ϕ*, and *M* in Equation 1 under least squared loss. *R*^2^ ranges from 0 to 1, with 1 indicating perfect sinusoidal wave fitting, and 0 indicating no fitting at all. Equivalently, we could re-write Equation 1 as

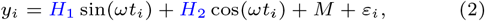

where *H*_1_ = *A* cos(*ωϕ*), and *H*_2_ = *A* sin(*ωϕ*), which turns into a linear regression problem.

### Power calculation

#### Analytical power calculation

According to linear model theories, the *F* statistics for the circadian model in Equation 1 can be derived as

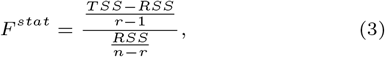

where *n* is number of independent samples, *r* = 3 is number of parameters (i.e., *A*, *ϕ*, and *M* in Equation 1).

Under the null hypothesis that there is no circadian rhythmicity. i.e., *A* = 0 in Equation 1, or equivalently, *H*_1_ = *H*_2_ = 0 in Equation 2,

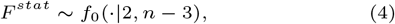

where 2 and *n* – 3 are the degrees of freedom of the *F* distribution, and *f*_0_ denotes a regular *F* distribution with non-centrality parameter 0 [42].

Under the alternative hypothesis that there exists a circadian rhythmicity pattern. i.e., *A* ≠ 0 in Equation 1, or equivalently, *H*_1_ ≠ 0 or *H*_2_ ≠ 0 in Equation 2, the *F* statistics follows a non-central *F* distribution,

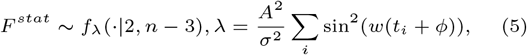

where 2 and *n* – 3 are the degrees of freedom of the *F* distribution, and *f*_λ_ denotes a non-central *F* distribution with non-centrality parameter λ. (see Supplementary Section 1 for proof).

Figure 2b shows the relationship between the null and alternative distributions. By assuming the type I error rate at the rejection boundary *F*\* is *α*, the relationship between *α* and the power 1 – *β* is

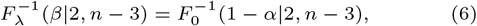

where *F*_λ_ (*x*|*df*_1_, *df*_2_) represents cumulative density function of *f*_λ_(·|*df*_1_, *df*_2_) evaluated at *x*, and *F*_0_ (*x*|*df*_1_, *df*_2_) represents cumulative density function of *f*_0_(·|*df*_1_, *df*_2_) evaluated at *x*.

As shown in Figure 2b, the non-centrality parameter λ controls the degree of separation of the null distribution *f*_0_ and the alternative distribution *f*_λ_. The larger the λ is, the more likely the alternative distribution will be away from the null distribution, and the higher power a gene will achieve. We thus define λ as the total effect size for the circadian power calculation. By inspecting the total effect size 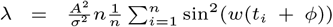, this non-centrality parameter can be decomposed into three parts: (i) sample size *n*, (ii) intrinsic effect size *r* = *A/σ* (closely relate to the goodness of fit statistics *R*^2^), and (iii) sampling design effect 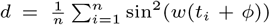. We then discuss the impact of each of these components on the circadian rhythmicity power calculation:

1. <u>Sample size</u>: As expected, given fixed *d* and *r*, a larger sample size *n* will result in a larger total effect size λ, and achieve a higher statistical power.
2. <u>Intrinsic effect size</u>: Intuitively, a larger circadian amplitude A with smaller residual variability *σ* will lead to a better sinusoidal curve fitting (i.e. larger *R*^2^). Our formula suggests that circadian fitting parameters *A* and *σ* work together as an intrinsic effect size *r* = *A/σ* and has a quadratic effect on the total effect size λ.
3. <u>Sampling design effect</u>: The sampling design effect 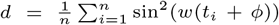 is more complicated, because it involves both observed *t_i_* and the unknown parameter phase shift *ϕ*. In general, given an arbitrary circadian sampling design, we need to estimate *ϕ* before performing power calculation. Fortunately, the power calculation for the evenly-spaced sampling design is independent of the phase value (i.e., phase-invariant). For example, [28] recommended a collection of 12 time points (every 2 hours) per cycle across 2 full cycles, which belongs to the evenly-spaced sampling design. Such active design is commonly seen in animal studies or human blood studies, where researchers can control the exact time to sacrifice the animal or to collect blood. The following theorem (phase-invariant property) shows that the sampling design effect *d* is a constant under the one-period one-sample evenly-spaced design, i.e., 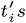 are evenly spread within one period with only one sample per time point.

##### Theorem 1

(Phase-invariant property - one-period one-sample) *Assuming there is a total of n ZT points t_i_*(1 ≤ *i* ≤ *n*) *within a circadian period* 2*π/ω, which are ordered such that t_i_* < *t*_i+1_ *for all* 1 ≤ *i* ≤ *n* – 1. *If n* ≥ 3, *and t_i_ is evenly-spaced over the period* (*i.e*., *t*_*i*+1_ – *t_i_* = *C for all* 1 ≤ *i* ≤ *n* – 1, *C* > 0 *is a fixed time interval*, (*t_i_* + 2*π/ω*) – *t_n_* = *C*), *then regardless of the value for ϕ, we have*

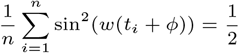

The proof is given in Supplementary Section 2. It can immediately be extended to the following corollary.

##### Corollary 1

(Phase-invariant property - multi-period multi-sample). *For multi-period (two or more cycles) multisample evenly-spaced design, the sampling design effect is phase-invariant*.

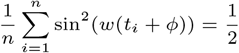

This is because the multi-period multi-sample evenly-spaced design just replicates the one-period one-sample evenly-spaced design, therefore the average of them remains to be 1/2.

#### Assumptions underlying the circadian modeling framework

The proposed circadian modeling framework has two underlying assumptions: (i) the relationship between the expression level of a gene and the ZT follows a sinusoidal wave curve; (ii) the error terms of each sample on top of the sinusoidal wave curve follows independent and identical Gaussian distribution. We discuss the implication of sinusoidal assumption on sampling design in Section 3.3 and demonstrate that the *F* statistics is robust against various types of violation of model assumptions in Section 3.4.

#### Alternative power calculation method by Monte-Carlo simulation

Without the proposed analytical method CircaPower, a conventional method for circadian detection power calculation is by Monte-Carlo simulation (MC), which assumes known A, *ϕ*, *M*, *σ* and *t_i_*, 1 ≤ *i* ≤ *n* in Equation 1. The detailed algorithm for MC is described as following:

1. Given the ZT 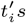 for *n* samples (1 ≤ *i* ≤ *n*) and key parameters (*A*, *ϕ*, *M*, and *σ*), we simulate gene expression *y_gi_* based on Equation 1, where 1 ≤ *g* ≤ *G* is the gene index and *G* is the total number of genes. *G* = 10, 100 genes is used in the simulation comparison between CircaPower and MC algorithm in Section 3.1.
2. We apply the cosinor method [31] to derive the rhythmic p-value p*_g_* for each gene *g*(1 ≤ *g* ≤ *G*). Given a pre-specified alpha level *α*, the power of MC algorithm is calculated as 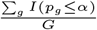.

Although both the CircaPower and the MC algorithm rely on the *F* statistics for rhythmicity detection, the CircaPower has several obvious advantages over the MC algorithm. First of all, the explicit representation of total effect size in CircaPower provides insights on the three determining factors (*n,r,d*) in circadian detection power calculation while it is hard for MC algorithm to determine selections and trends on the many parameters (*A*, *ϕ*, *M*, *σ*, and *t_i_*, 1 ≤ *i* ≤ *n*). In addition, our simulation shows the closed-form solution by CircaPower is at least 10,000 folds faster than the MC algorithm (see Section 3.1). More importantly, even though both approaches can calculate power given sample size, only CircaPower can directly solve the inverse problem of deriving the smallest sample size meeting the desired detection power, while MC algorithm needs repeated interpolation to obtain an answer.

## Simulation

Throughout simulation and real application, we control type I error *α* = 0.001 for circadian power calculations to account for potential multiple comparisons.

### Comparison between CircaPower and the MC algorithm

We compare CircaPower with the MC algorithm described in Section 2.2.3. For both methods, the ZT points are simulated from one-period one-sample evenly-spaced design for the ease of discussion, which enjoys the phase-invariant property (*d* = 1/2, Theory 1). Since phase shift *ϕ* and MESOR *M* has no impact on circadian detection power calculation in this case, we fix *ϕ* = 0 and *M* = 10. We evaluate their power derived at a grid of *A* = (0.4, 0.8, 1, 1.2) and *σ* = (1, 2, 3, 4). Note that for CircaPower, we only need the underlying parameters *A/σ*, *t_i_*, and *ϕ* to perform power calculation, which does not rely on the simulated dataset. We simulate the data for the purpose of evaluating the MC algorithm.

Figure S1a shows that the power calculated from CircaPower is almost identical to the MC algorithm, corroborating the correctness of the closed-form solution in CircaPower. In addition, the power increases with respect to (i) larger *n*, (ii) larger A, and (iii) smaller *σ*, which are consistent with our theoretical formula of the total effect size λ.

As discussed in Section 2.2.1, the two curve fitting parameters *A* and *σ* work together as the intrinsic effect size r = A/σ and therefore can be reduced to one parameter in MC approach. We further validate this observation by co-varying *A* and *σ* simultaneously (i.e., *A* = *σ* = 1, 2, 3) while keeping their ratio as a constant (i.e., *r* = 1). Figure S1b shows that the power trajectories remain the same, indicating the proposed intrinsic effect size is sufficient to capture the goodness-of-fit of the model for the power calculation.

In terms of computing time, to generate all the results in Figure S1a, it takes 1.84 seconds for the CircaPower using 1 CPU thread on a regular PC (8th Gen Intel Core i5-8250U Quad-Core processor, 1.60 GHz), while it requires 8 hours for the MC algorithm using the same computing resource. With parallel computing, the computing time reduces to 0.13 seconds for CircaPower using 40 CPU threads on a Linux server (Intel Xeon Gold 6130, 2.10GHz), while it still needs 24 minutes for the MC algorithm.

### Impact of sampling design on CircaPower

Since the sample collection scheme for active design and passive design are quite different, we will discuss them separately. For active designs, we vary the intrinsic effect *r* = 0.8, 1, 1.5, 2 and *n* = 12, 24, 36, 48. For passive designs, we vary the intrinsic effect *r* = 0.4, 0.8, 1, 1.2 and *n* = 12, 24, …, 180. The maximal sample size we use for the active design is smaller than that of the passive design, because the estimated intrinsic effect is usually higher in animal studies compared with human studies (see Table 1). Due to the fact that not all designs have the phase-invariant property, we also vary the phase shift *ϕ* = 0, 3, 6.

**Table 1.**
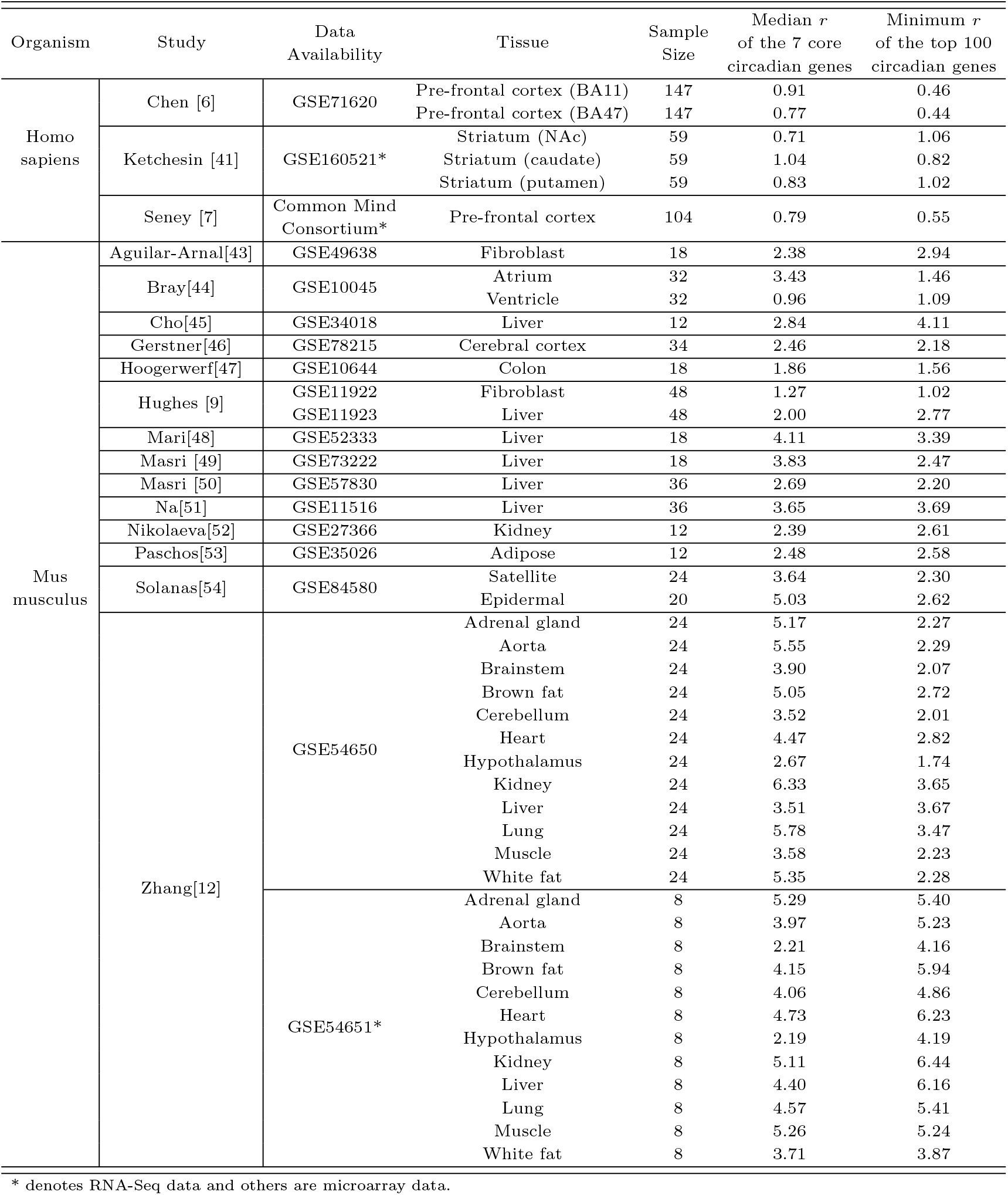
Intrinsic effect sizes for public available transcriptomic circadian data, including 3 passively designed human postmortem brain studies and 14 actively designed mouse studies from 20 types of tissues. These data are processed using the cosinor method [31]. Two types intrinsic effect sizes are used: (i) median *r* of the 7 core circadian genes; (ii) minimum *r* of the top 100 significant circadian genes. These intrinsic effect sizes can be used as a reference resource when investigators need to perform power calculation without any pilot data.

For a typical active design, researchers usually need to control the number of (i) ZT points per cycle; (ii) replicates at each time point within a cycle; and (iii) cycles. Because of the periodicity property of the sinusoidal curve in the cosinor model, (ii) and (iii) are statistically equivalent. Therefore, for the ease of discussion, we summarize the following two key parameters for an active design: (i) number of ZT points per cycle *N_T_*; (ii) total number of samples *n*. The number of replicated samples (at the same ZT across all cycles) could be calculated as *n/N_T_*.

We denote the active design scheme with *N_T_* points per cycle as FixTimeNT an the one-period one-sample evenly-spaced design (i.e., FixTime-*n*) as the EvenSpace. For *N_T_* = 3, 4, 6, *n*, Figure 3a shows that (i) the power curves are the same regardless of *phi*, confirming the phase-invariant property; (ii) the power trajectories for different 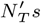 are also identical, which implies that under evenly-spaced sampling design with *N_T_* ≥ 3, the detection power only depends on the total number of samples *n* but not the *N_T_*. Note that these arguments are purely based on the statistical power given sinusoidal wave assumption. In reality, less number of time points may result in unstable circadian curve fitting (see Section 3.3).

**Fig. 3.**
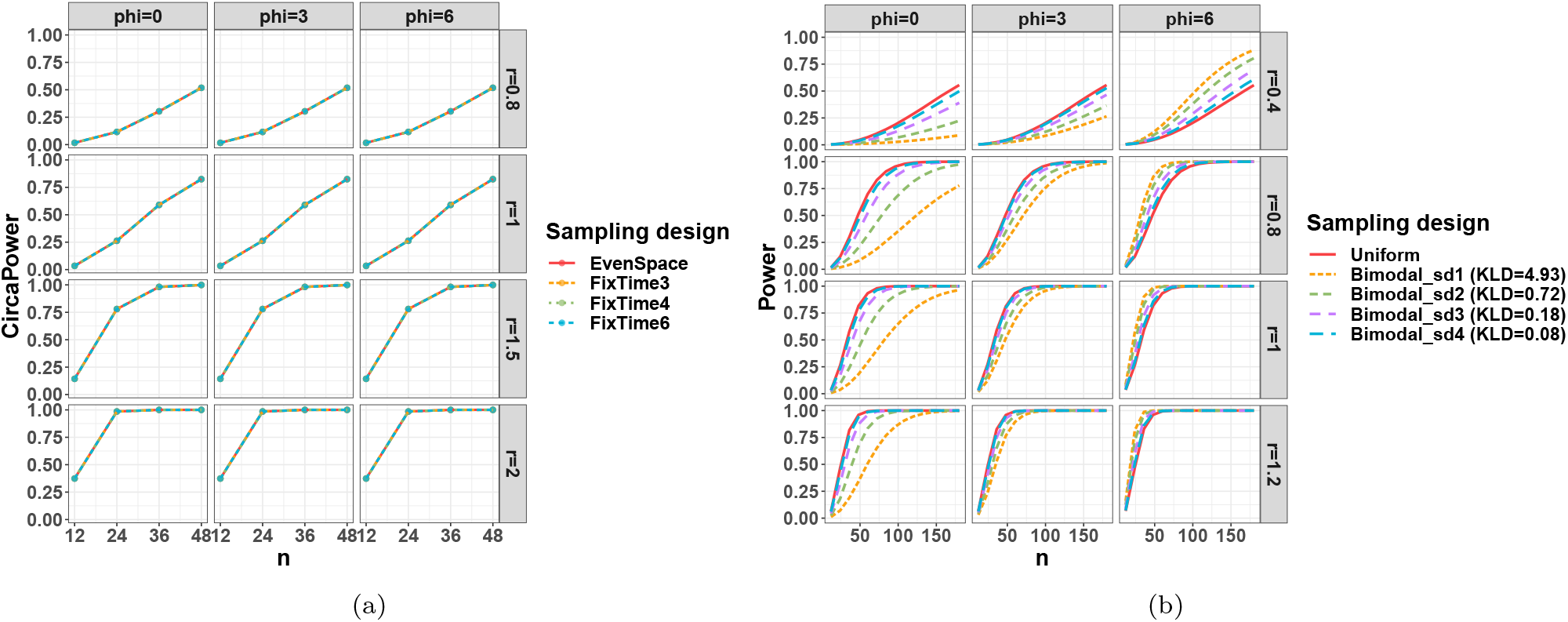
Sampling design effect on circadian power calculation. (a) shows the sampling design effect for active design; (b) shows the sampling design effect for passive design.

For passive designs, the collection of the ZT cannot be controlled. We therefore simulate 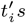 from (i) uniform distribution (uniform design): 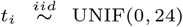; and (ii) bimodal Gaussian distributions (bimodal designs): *t_i_*~*p_i_N*(7,*sd*) + (1 – *p_i_*)*N*(17, *sd*); *p_i_* ~ Bernulli(0.5). We allow *sd* = 1, 2, 3, 4 and calculate their corresponding Kullback–Leibler divergence (KLD) against the uniform distribution as a relative measurement to benchmark their divergence from the uniform. Figure 3b shows that for the uniform design, the power trajectory is close to phase-invariant. This is expected since the uniform distribution is a random realization of the evenly-spaced sampling design, and the impact of phase on the individual *t_i_* will average out. The bimodal designs show phase-dependent circadian power trajectories and the impact of phase influence increases as the distribution deviates more from uniform (i.e., larger KLD). Specifically, the power loss of bimodal designs when *ϕ* = 0, 3 is significant when KLD is 0.72 or greater (green and yellow curve) while negligible when KLD is only 0.18 or smaller (purple and blue curve). In fact, since the phase shift impacts the sampling design through 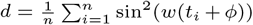, it achieves higher power if the mode of the ZT distribution occurs at the underlying peak/trough time. In real omics applications, circadian genes usually have different phase shift values over the day. If the collected ZT distribution is far away from the uniform distribution, the detection power of each circadian gene would be affected differently across the genome as a result of its unique phase shift.

### Impact of the number of time points per cycle on CircaPower

From the perspective of circadian power calculation, Figure 3a implies that the evenly-spaced sampling design is phase-invariant as long as *N_T_* ≥ 3, which is further corroborated by Corollary 1. However, in the perspective of curve fitting, smaller number of time points may not necessarily guarantee the goodness-of-fit for a sinusoidal curve, resulting in potentially false positive findings.

To explore the impact of number of *N_T_* on the goodness-of-fit for a sinusoidal wave, we simulate expression data from the sinusoidal model and perform non-parametric curve fitting to identify the minimum *N_T_* necessary to capture the sinusoidal wave curve (see detailed simulation setting in Supplementary Section 4). The data points and fitted smooth curves in one circadian cycle [0, 24] are shown in Figure S2. When *N_T_* ≤ 4, it is uncertain whether the underlying curve fitting is a sinusoidal wave. Only when *N_T_* increases to 6 or more, the curve fitting is stable and almost identical to the underlying sinusoidal wave. To further justify this choice, we calculate the root mean square distance between the fitted non-parametric curve and the underlying sinusoidal curve at evenly spaced 1000 points between [0, 24] and plot against *N_T_*. The elbow plot (Figure S3) suggests that *N_T_* = 6 is an inflection point after which the change of distance between the fitted curve and the underlying curve becomes stably small. Therefore, considering both circadian power calculation and smooth curve fitting, our results suggest *N_T_* = 6 to be the minimum number of ZT points to fully capture the circadian rhythmicity pattern, which is commonly adopted in the literature.

### Robustness analysis of *F* statistics

To examine the robustness of our method when the *iid* Gaussian assumption is violated, we investigate the type I error control of *F* statistics in the following scenarios: (i) heavy tail error distribution (i.e., student t distribution); (ii) existence of outliers; (iii) non-independent Gaussian errors. For all these simulations, *G* = 10, 000 noisy genes are simulated with error term *ε_gi_*’s specified above. By declaring circadian rhythmicity at 5% nominal *α* level, we will evaluate the actual type I error rate of the *F* test from the cosinor model. Since CircaPower is built on the *F* statistics for rhythmicity detection it will be benchmarked as robust if the actual type I error rate is close to the nominal *α* level. The detailed simulateion setting is provided in Supplementary Section 3. Since our goal is to evaluate the type I error rate control, which does not involve in any multiple testing issue, we directly use 5% nominal *α* level. Figure S4 shows that the cosinor model only has slightly conservative the type I error from cosinor model when error terms are drawn from very heavy tail t distribution (df = 2.5 or 3) while maintains accurate type I error rate in all other scenarios suggesting the robustness against the three types of violation in general.

## Real application

### CircaPower for human studies with passive design

We investigate the power trajectories of human studies using three human post-mortem brain transcriptomic studies (Chen [6], Seney [7] and Ketchesin [41]) with different time of death distribution. Detailed descriptions of each dataset can be found in the original papers. Briefly, Chen [6] and Seney [7] performed gene expression circadian analysis using microarray (*n* = 147) and RNA-seq (*n* = 104) respectively using pre-frontal cortex tissues; and Ketchesin [41] performed RNA-seq gene expression circadian analysis with *n* = 59 participants using dorsal and ventral striatum tissues.

To estimate the intrinsic effect sizes from the three brain studies, we apply the cosinor method [31] to identify genes with rhythmic patterns and obtain estimates for their amplitude 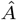 and noise level 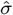. We estimate the intrinsic effect sizes 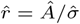 using the 7 core circadian genes, or the top 100 significant rhythmic genes (ranked by p-values from the cosinor method). The 7 core circadian genes include *Arntl, Dbp, Nr1d1, Nr1d2, Per1, Per2*, and *Per3*, which showed persistent circadian pattern across 12 mouse tissues [12]. The Homo sapiens section of Table 1 shows the estimated intrinsic effect sizes for: (i) median *r* of the 7 core circadian genes; (ii) minimum *r* of the 100 most significant circadian genes. The estimated intrinsic effect sizes for these three human studies range between 0.44 and 1.06 (see Table 1).

To demonstrate the power trajectories in real data, we vary intrinsic effect sizes *r* = 0.4, 0.6, 0.8, 1, which roughly cover the estimated range of the intrinsic effect sizes in the postmortem brain studies. The ZT points are sampled 1000 times from the kernel density estimated from the observed time-of-death distributions in these three studies (see top panels of Figure 4a). Since investigators in these human post-mortem brain studies have no control of sample collection time (i.e., time of death) and can only accept passive sampling design, the detection power curves are not phase-invariant. We vary phase shift *ϕ* = 0, 1, 2, 3, …, 12 and use a confidence band to represent the range of power achieved across phase shifts (see bottom panels of Figure 4a). For each scenario (i.e., fixed r, *n* and *ϕ*), the mean power among the 1000 times repetitions is reported. The power trajectory from an evenly-spaced design is also calculated as a comparison.

**Fig. 4.**
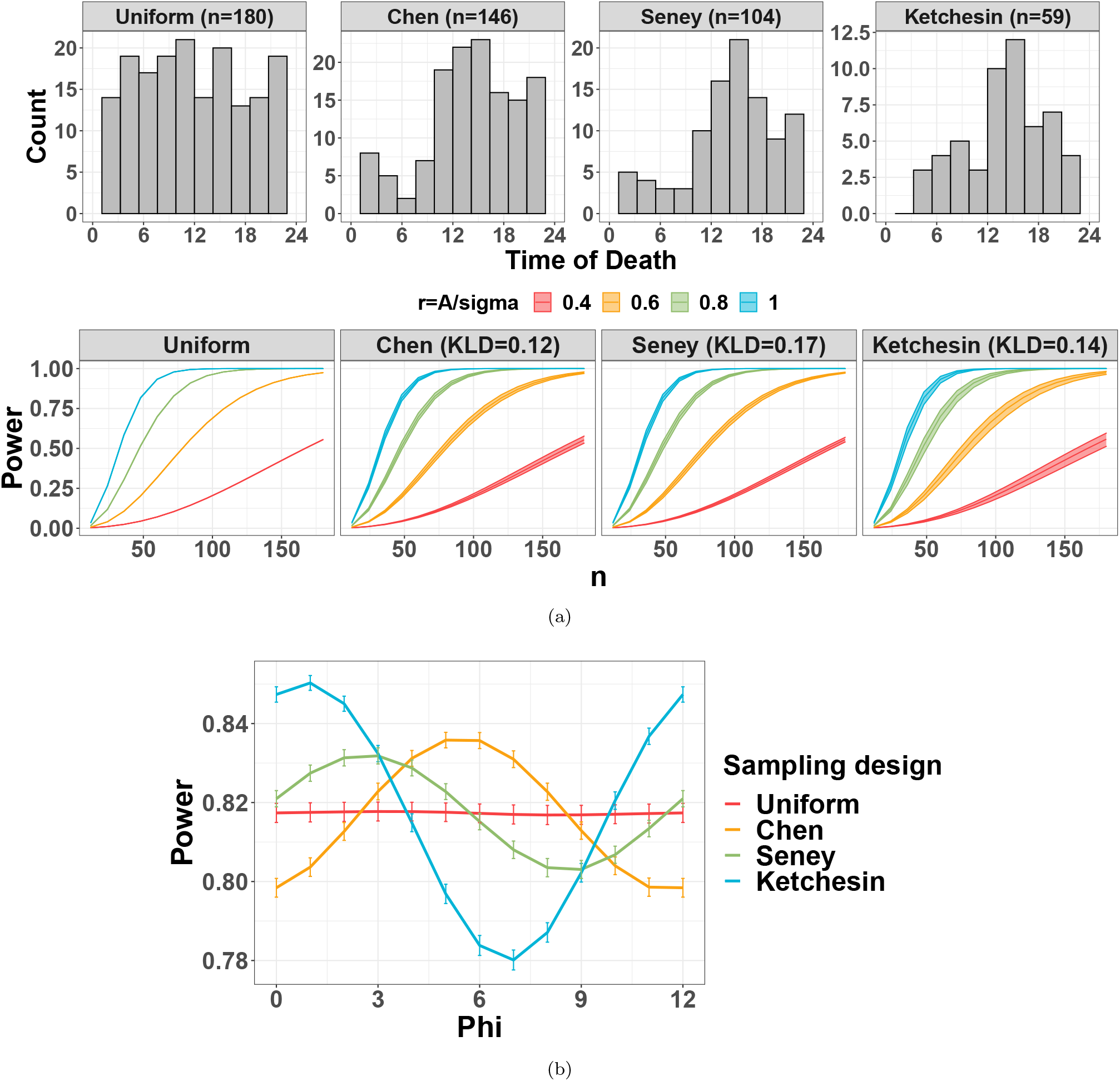
(a) demonstrates circadian power calculation using publicly available human datasets. The top panel shows the time of death distribution for Chen, Seney, and Ketchesin. The bottom panel shows the mean power trajectories of different study designs over 1000 repetitions with different intrinsic effect sizes *r* = (0.4, 0.6, 0.8, 1). The confidence bands represent the range of power achieved across phase values at *ϕ* = 0, 1, 2, 3, …, 12 for each scenario. (b) shows mean power trajectories across different *ϕ* when *n* = 120 and *r* = 0.6. For each *ϕ* and sampling distribution, we draw sampling times 1000 times and calculate corresponding power. Vertical bars indicate the 95% confidence interval of power estimates calculated form 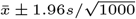 where 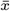 and *s* are mean and standard deviation of power estimates respectively. Maximum power drop (calculated by 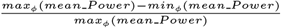 is 3.8%, 4.4%, and 9.9% respectively.

As expected, larger sample size *n* and larger intrinsic effect size *r* lead to a larger circadian power. In addition, the power trajectory is phase-invariant for the evenly-spaced sampling design with 0 band width, but not for the passive sampling designs from the three human studies. As discussed in Section 3.2, the power will depend on the relationship between the mode of the ZT distribution and the underlying peak/trough time. To further demonstrate the impact of phase on the non-uniform ZT distribution, we fix *n* = 120, *r* = 0.6, while varying *ϕ* = 0, 1, 2, 3, …, 12. As shown in Figure 4b, the power trajectories fluctuate across different *ϕ*’s when samples are drawn from irregular distributions in the three post-mortem studies, while the trajectory stays the same for evenly-spaced ZT. However, since the KLDs of the kernel densities estimated from the Chen, Seney and Ketchesin are relatively low (i.e., 0.12, 0.17, 0.14) compared with the bimodal designs in Section 3.2, the variation of power as a result of phase shift is small, with 3.8%, 4.4%, and 9.9% maximum drop, respectively.

### CircaPower for animal studies with with active design

We next examine the power trajectories of actively designed mouse studies using 14 mouse gene expression circadian data [43, 44, 45, 46, 47, 9, 48, 49, 50, 51, 52, 53, 54, 12] from 20 types of tissues. Sample sizes of each study tissue are shown in Table 1. To estimate the intrinsic effect sizes of these tissues, we apply the cosinor method [31] similarly to identify genes with rhythmic patterns and obtained estimates for their amplitude 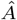 and noise level 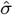. The estimated intrinsic effect sizes for the median *r* of the 7 core circadian genes and the minimum *r* of the top 100 significant circadian genes are shown in the Mus musculus section of Table 1, ranging from 0.96 to 6.33, a much larger magnitude than previous human studies. This is reasonable since human studies are usually more heterogeneous in terms of genetics and environmental background. We thus fix the intrinsic effect sizes to be *r* = 1, 2, 3, 4 in our subsequent power calculation. Since these experiments employ an evenly-spaced active sampling design, the sampling design factor is a constant (i.e., *d* = 1/2, Corollary 1) regardless of the phase value.. As a result, we employ the one-sample one-period evenly-spaced design (see left panel of Figure 5) for the purpose of power calculation. By further assuming the alpha levels to be *α* = 0.05, 0.01, 0.001, the power trajectories with respect to sample size *n* is shown in Figure 5 (right panel).

**Fig. 5.**
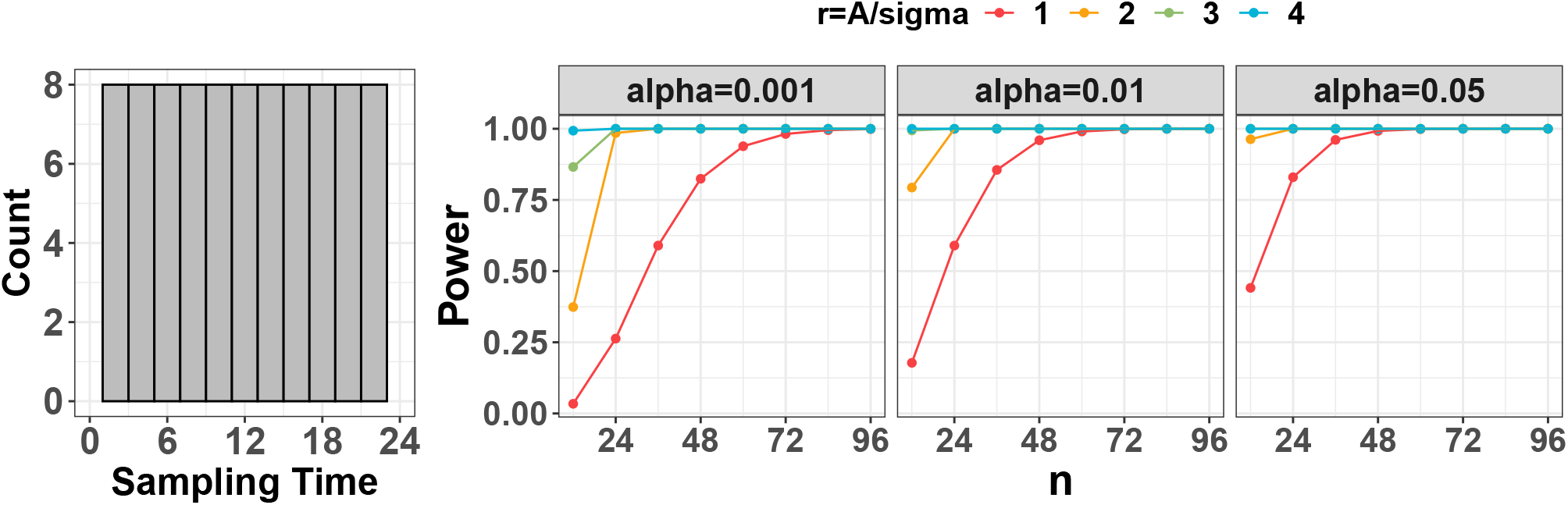
Circadian power calculation using publicly available mouse datasets. The left panel shows the distribution of one-sample one-period evenly-spaced design. The right panel shows the power trajectories for each of the type I error control *α* = (0.001, 0.01, 0.05) assuming intrinsic effect sizes *r* = (1, 2, 3, 4).

### Case study: circadian power calculation using mouse pilot dataset

To demonstrate how to perform circadian power calculation using pilot dataset from scratch, we utilize a circadian gene expression data in mouse with skeletal muscle, which is part of the mouse pan-tissue gene expression circadian microarray data [12]. Detailed description of this dataset has been described previously [8]. Briefly, 24 mouse muscle samples were collected (every 2 hours) across 2 full cycles. With this pilot data, we perform genome-wide circadian rhythmicity detection using the cosinor method [31]. Under *p* < 0.001, we identify 716 significant genes showing circadian pattern. We similarly estimate the intrinsic effect sizes for: (i) median *r* of the 7 core circadian genes; (ii) minimum *r* of the top 100 significant circadian genes. The resulting intrinsic effect sizes are 3.58 and 2.23 respectively. By assuming different *α* to be 0.05, 0.01, 0.001, the power curves with respect to sample size are shown in Figure S5. We observe that *n* = 12 can achieve 97.1% and 50.5% detection power for the two intrinsic effect sizes at *α* = 0.001.

## Discussion

In this paper, we propose an analytical framework, CircaPower, to calculate the statistical power for circadian gene detection. To the best of our knowledge, this is the first analytical method to perform circadian power analysis. In simulations, we not only demonstrate the CircaPower is fast and accurate, but also show that its underlying cosinor model is robust against violations of model assumptions. In real applications, we show the performance of the CircaPower in mouse studies and postmortem human studies. In addition, we obtain the estimated intrinsic effect sizes from publicly available human and mouse transcriptomic circadian data. These summarized intrinsic effect sizes can be used as a reference resource to facilitate investigators without pilot data to perform circadain power calculation. In case study, we also demonstrate circadian power calculation step-by-step given a pilot dataset.

Our method has several advantages. To begin with, the theoretical framework suggests that the power calculation is related to a total effect size, which can be decomposed into sample size, intrinsic effect size (representing goodness-of-fit of circadian curve), and sampling design factor. Moreover, the sampling design factor brings about the concept of active design and passive design when samples are collected. This is an important concept in circadian experiment design, since the ZT collection for human (passive design) and animal (active design) could be quite different. After that, we demonstrate the phase-invariant property of the evenly-spaced sampling design, which provides theoretical foundation for the design of many published circadian studies. In addition, the closed-form formula in CircaPower allows unique inverse calculation of sample size given desired power at fast computing speed compared with the conventional MC algorithm. In this paper, we also systematically examine the intrinsic effect sizes of published mouse or human gene expression circadian data, which could provide guidance for future researchers to design their transcriptomic circadian experiment when pilot data are not available. Although we present our work using transcriptomic data, CircaPower is applicable to other omics data, such as DNA methylation, ChIP-Seq proteomics, metabolomics, and clinical data (e.g., body temperature).

Our work has the following limitations and future work. Firstly, the CircaPower assumes a pre-fixed alpha level. We intentionally select a more stringent alpha (e.g., *α* = 0.001) to account for multiple comparison when thousands of genes are tested simultaneously. Additional modeling is needed to extend for calculating genome-wide power calculation while controlling false discovery rate. Secondly, in addition to detecting genes with rhythmic pattern, another important research question is to identify differential circadian pattern [55, 56, 57, 32] (i.e., the circadian pattern is disrupted because of the treatment or condition), which will be another future direction. Thirdly, the Gaussian assumptions are widely assumed in biomedical research, and we have demonstrated that the cosinor model is robust against violations of Gaussian assumptions. If an investigator still worries about these assumptions, we would recommend data transformations (e.g. Box–Cox transformation) before applying our method. Our previous work [32] has shown that the Box-Cox transformation can rescue the normality assumption for circadian rhythmicity detection using cosinor models. Similar justifications have been adopted in the literature. For example, though the student t test also assumes Gaussian assumptions, but it is still widely used in the literature, as long as there are methods to rescue the violation (i.e., data transformation). Lastly, as discussed in the introduction, both parametric and non-parametric models are popular and widely used in the literature. In the proposal, we only focus on the cosinor model for its simplicity and accurate statistical inference [32]. Further extending the current framework to a more flexible family of circadian pattern is of biological interests to the general circadian research field.

To allow easy application by other researchers, our methods have been implemented in the R package CircaPower, which is publicly available in github (https://github.com/circaPower/CircaPower).

## Data Availability

Seney [7] dataset is available in the Common Mind Consortium https://www.nimhgenetics.org/available_data/commonmind/ through an approval process. All other datasets are publicly available on the NCBI GEO database with accession numbers shown in Table 1.

## Acknowledgments

We thank the anonymous reviewers for their valuable suggestions.

## Key points

- The circadian clock controls oscillations in a wide variety of physiological processes, and the disruption in clock and circadian gene expression was found to be linked to many diseases. Though recent transcriptomic studies have been successful in revealing the circadian rhythmicity in gene expression, the power calculation for omics circadian analysis have not been fully explored. To our knowledge, we are the first to develop rigorous statistical methods for circadian power calculations.
- It is unclear what factors will impact the power calculation of circadian analyses. Our theoretical framework is the first to determine three key factors in circadian power calculation, including sample size, intrinsic effect size and sampling design factor.
- We further summarize and document the intrinsic effect sizes from 3 human postmortem brain studies and 14 mouse studies from 20 types of tissues, which would facilitate researchers without pilot data to perform circadian power calculation.
- Our method CircaPower has been implemented in an R package, which is made publicly available on GitHub (https://github.com/circaPower/CircaPower).

**Wei Zong** is a PhD student in the Department of Biostatistics at University of Pittsburgh. She is interested in statistical modelling on high-dimensional genomics data and molecular circadian rhythmicity analysis.

**Dr. Marianne L. Seney** is Assistant Professor of Psychiatry at University of Pittsburgh. Her lab focuses research on the sex differences in psychiatric disorders, dendritic spine/microglia alterations and molecular rhythm disruptions in brain disorders.

**Dr. Kyle D. Ketchesin** is Assistant Professor of Psychiatry at University of Pittsburgh. He is interested in investigating the epigenetic mechanisms underlying circadian dysfunctions in mood disorders, particularly depression.

**Michael T. Gorczyca** is a Ph.D. Student in the Department of Computational and Systems Biology at University of Pittsburgh. His research interest includes developing methodology to quantify circadian dysfunction and find associations between circadian dysfunction and psychiatric disorders.

**Dr. Andrew C. Liu** is Associate Professor in Department of Physiology Functional Genomics at University of Florida College of Medicine. The major focus of his lab is to study the molecular, cellular, and physiological mechanisms of circadian (~ 24) clocks in mammals.

**Dr. Karyn A. Esser** is Professor in Department of Physiology Functional Genomics at University of Florida College of Medicine. Her lab focuses research on the role of circadian rhythms and the molecular clock mechanism in skeletal muscle homeostasis and health.

**Dr. George C. Tseng** is Professor in the Department of Biostatistics at University of Pittsburgh. His research group focuses on developing rigorous, timely and impactful methodologies in the area of genomics and bioinformatics, to help understand disease mechanisms and improve disease diagnosis and treatment.

**Dr. Colleen A. McClung** is Professor of Psychiatry and Clinical and Translational Science at University of Pittsburgh. She is interested in studying the association between circadian clock and various psychiatric disorders, including bipolar disorder, major depression and drug addiction.

**Dr. Zhiguang Huo** is Assistant Professor in the Department of Biostatistics at University of Florida. His research interest includes developing statistical and machine learning methodology for the broad field of genomics and bioinformatics.

