## Supplementary Materials for "Experimental Design and Power Calculation in Omics Circadian Rhythmicity Detection"

### S1 Proof of $F$ statistics distribution under the alternative

#### S1.1 Derivation of $F$ statistics

Denote $y_i$ as the expression value of one gene for sample $i (1 \leq i \leq n)$, where $n$ is the total number of samples. $t_i$ is the Zeitgeber time (ZT) for sample $i$. We assume the gene expression value is a sinusoidal wave function of the ZT plus some random noises:

$$y_i = A \sin(\omega(t_i + \phi)) + M + \varepsilon_i, \tag{1}$$

where $\varepsilon_i$ is the error term for sample $i$; we assume $\varepsilon_i$'s are identically and independently distributed (i.e., *iid*) and $\varepsilon_i \sim N(0, \sigma^2)$, where $\sigma$ is the noise level. To benchmark the goodness of sinusoidal wave fitting, we define the coefficient of determination

$$R^2 = 1 - \frac{RSS}{TSS}$$

where $RSS = \sum_{i=1}^{n}(y_i - \hat{y}_i)^2$, $TSS = \sum_{i=1}^{n}(y_i - \bar{y})^2$, $\hat{y}_i = \hat{A}\sin(\omega(t_i + \hat{\phi})) + \hat{M}$, $\bar{y} = \sum_i y_i/n$, with $\hat{A}$, $\hat{\phi}$, and $\hat{M}$ being the fitted value for $A$, $\phi$, and $M$ in Equation 1 under least square loss. $R^2$ ranges from 0 to 1, with 1 indicating perfect sinusoidal wave fitting, and 0 indicating no fitting at all. Denote $H_1 = A\cos(\omega\phi)$, $H_2 = A\sin(\omega\phi)$. Equivalently, we could re-write Equation 1 as

$$y_i = H_1 \sin(wt_i) + H_2 \cos(wt_i) + \varepsilon_i \tag{2}$$

Under the linear model framework, the $F$ statistics ($F^{stat}$) could be derived as:

$$F^{stat} = \frac{\frac{TSS - RSS}{r-1}}{\frac{RSS}{n-r}}$$

#### S1.2 Derivation of $F$ statistics distribution under the null and the alternative

**Lemma S1.1.** *Under the null distribution in the linear model framework, $F^{stat} \sim f_0(\cdot|r-1, n-r)$, where $f_0(\cdot|r-1, n-r)$ denotes a regular F distribution with degrees of freedom $r-1$ and $n-r$.*

*Proof.* The proof is given in [3]. □

**Lemma S1.2.** *If $X \sim N_p(\mu, \Sigma)$ and $\Sigma$ is positive definite variance-covariance matrix for the $p$ dimensional multivariate normal distribution, then*

$$X^\top \Sigma^{-1} X \sim \chi^2_p(\lambda),$$

where $\lambda = \mu^\top \Sigma^{-1} \mu$ is the non-centrality parameter for the $\chi^2_p$ distribution with degree of freedom $p$.

*Proof.* The proof is given in [3]. □

**Lemma S1.3.** *In a linear model framework, if the design matrix $X \in \mathbb{R}^{n \times p}$, the regression coefficient $\beta \in \mathbb{R}^p$, there are $q$ hypotheses to be tested $A^\top \beta = b$, where $A \in \mathbb{R}^{p \times q}$, $b \in \mathbb{R}^q$ represents the true parameters $b = A^\top \beta^*$.*

*We have the following results:*

$$F^{stat} = \frac{(A^\top \hat{\beta})^\top B^{-1} (A^\top \hat{\beta})}{RSS} = \chi^2_q(\lambda)/\chi^2_{n-r} \sim f_\lambda(\cdot|q, n-r),$$

*where $f_\lambda(\cdot|q, n-r)$ denotes a non-central F distribution with non-centrality parameter $\lambda = (A^\top\beta)^\top B^{-1}(A^\top\beta)$ and degrees of freedom $q$ and $n-r$; $B = A^\top(X^\top X)^g A$.*

*Proof.*

$$A^\top \hat{\beta} = A^\top (X^\top X)^g X^\top Y$$

$$A^\top \hat{\beta} \sim N_q((A^\top \beta), \sigma^2 A^\top (X^\top X)^g A)$$

If we denote $B = A^\top (X^\top X)^g A$, and base on Lemma S1.2, we have:

$$\frac{(A^\top \hat{\beta})^\top B^{-1} (A^\top \hat{\beta})}{\sigma^2} \sim \chi^2_q(\lambda)$$

where $\lambda = (A^\top \beta)^\top B^{-1} (A^\top \beta)$. In addition, since

$$\frac{RSS}{\sigma^2} = \frac{Y^\top (I - P) Y}{\sigma^2} \sim \chi^2_{n-r},$$

therefore we could derived the following relationship [4]:

$$F^{stat} = \frac{(A^\top \hat{\beta})^\top B^{-1} (A^\top \hat{\beta})}{RSS} = \chi^2_q(\lambda)/\chi^2_{n-r} \sim f_\lambda(\cdot|q, n-r)$$

□

**Theorem S1.4.** *Under the sinusoidal model assumption (Equation 2), under the alternative hypothesis (there is a circadian fitting, i.e., $H_1 \neq 0$ or $H_2 \neq 0$), we have*

$$F^{stat} \sim f_\lambda(\cdot|2, n-3),$$

*where* $\lambda = \frac{A^2}{\sigma^2} \sum_i \sin^2(w(t_i + \phi))$.

*Proof.* Fitting the sinusoidal model Equation 2 into the linear model framework, the design matrix is

$$X = \begin{pmatrix} \sin(\omega t_1) & \cos(\omega t_1) \\ \cdots & \cdots \\ \sin(\omega t_n) & \cos(\omega t_n) \end{pmatrix}$$

And

$$X^\top X = \begin{pmatrix} \sum_i \sin^2(\omega t_i) & \sum_i \sin(\omega t_i)\cos(\omega t_i) \\ \sum_i \sin(\omega t_i)\cos(\omega t_i) & \sum_i \cos^2(\omega t_i) \end{pmatrix}$$

The regression coefficient is

$$\beta = \begin{pmatrix} H_1 \\ H_2 \end{pmatrix}$$

The joint hypotheses are $H_1 = 0$ and $H_2 = 0$, which is equivalent to the following:

$$\begin{pmatrix} 1 & 0 \\ 0 & 1 \end{pmatrix} \begin{pmatrix} H_1 \\ H_2 \end{pmatrix} = \begin{pmatrix} 0 \\ 0 \end{pmatrix}$$

By denoting $A = I_2$, the hypothesis is equivalent to $A\beta = 0$. According to Lemma S1.3, we have

$$F^{stat} \sim f_\lambda(\cdot|2, n-3),$$

$$\begin{aligned} \lambda &= \frac{\beta^\top X^\top X \beta}{\sigma^2} \\ &= \frac{A^2}{\sigma^2} \sum_i \sin^2(w(t_i + \phi)) \end{aligned}$$

□

#### S2 Proof of the phase-invariant property in the one-sample one-period evenly-spaced design

**Lemma S2.1.** *For* $n \in \mathbb{N}^+$, $a \in \mathbb{R}$, $d \in \mathbb{R}$, *define* $R = \frac{\sin(Nd/2)}{\sin(d/2)}$.

$$\sum_{i=0}^{N-1} \cos(a + id) = \begin{cases} n\cos(a) & \text{if } \sin(d/2) = 0 \\ R\cos(a + \frac{1}{2}(n-1)d) & \text{otherwise} \end{cases}$$

*Proof.* See [2]. □

**Corollary S2.1.1.** *When* $n \geq 3$, $\phi \in \mathbb{R}$*;*

$$\frac{1}{n}\sum_{i=0}^{n-1} \cos(\frac{4\pi i}{n} + \phi) = 0$$

*With that being said, the evenly spaced design with* $n \geq 3$ *time points will achieve the same design effect, regardless of phase* $\phi$

*Proof.* Following Lemma S2.1, set $N = n$, where $n \geq 3$, $d = 4\pi/n$, then $\sin(d/2) = \sin(2\pi/n) \neq 0$, $R = 0$.

$$\begin{aligned}\frac{1}{n}\sum_{i=0}^{n-1} \cos(\frac{4\pi i}{n} + \phi) &= R\cos(a + \frac{1}{2}(N-1)d) \\ &= 0\end{aligned}$$

□

**Theorem S2.2** (Phase-invariant property - one-period one-sample)**.** *Assuming there is a total of n ZT points within a circadian period* $2\pi/\omega$, *the ZT* $t_i'$*s are ordered such that* $t_i < t_{i+1}$ *for all* $1 \leq i \leq n-1$. *If* $n \geq 3$, *and* $t_i$ *is evenly-spaced over the period (i.e.,* $t_{i+1} - t_i = C$ *for all* $1 \leq i \leq n-1$, $C > 0$ *is a fixed time interval,* $(t_1 + 2\pi/\omega) - t_n = C$*), then regardless of the value for* $\phi$,

$$\frac{1}{n}\sum_{i=1}^{n} \sin^2(w(t_i + \phi)) = \frac{1}{2}$$

*Proof.*

$$\begin{aligned}\frac{1}{n}\sum_{i=0}^{n-1} \sin^2(w(t_i + \phi)) &= \frac{1}{2n}\sum_{i=0}^{n-1}(1 - \cos(2w(t_i + \phi))) \\ &= \frac{1}{2} - \frac{1}{2n}\sum_{i=0}^{n-1}\cos(2\omega(\frac{2i\pi}{\omega n} + \phi))) \\ &= \frac{1}{2} - \frac{1}{2n}\sum_{i=0}^{n-1}\cos(\frac{4i\pi}{n} + \phi))) \\ &= \frac{1}{2}\end{aligned}$$

□

### S3 Simulation setting to evaluate the type I error control of F test in cosinor model

We consider three types of violation against the iid Gaussian assumption:

1. Heavy tail error distribution. Instead of sampling the error term $\varepsilon_{gi} \overset{iid}{\sim} N(0, \sigma^2)$, $1 \leq g \leq G$ and $1 \leq i \leq n$, we sample $\varepsilon_{gi} \overset{iid}{\sim} t(df)$, where $t(df)$ is the student t distribution with degree of freedom $df$. In general, the smaller the $df$ is, the heavier tail the error distribution is. When $df = 1$, the error distribution becomes the Cauchy distribution, and when $df \to \infty$, the error distribution converges to standard Gaussian distribution (i.e., $N(0, \sigma^2)$). To evaluate the impact of heavy tail error distribution on CircaPower, we simulate a grid of $df = (2.5, 3, 5, \infty)$. Figure S4a shows that when there is no or mild violation of the Gaussian assumption (i.e., $df = \infty$ or $df = 5$), the cosinor method achieves accurate type I error control (i.e., 5%). When there is moderate to severe violation of the Gaussian assumption (i.e., $df = 3$ or $df = 2.5$), type I error rate is only slightly conservative (i.e., below the nominal $\alpha = 0.05$). Putting together, the type I error rate of the cosinor method can be correctly controlled against the heavy tail error distribution.

2. Existence of outliers To evaluate the impact of outliers on type I error control, we replace $q$% of the expression values with outliers, where $q = (5, 10, 20)$. To be specific, for a gene $g$, there is $q$% chance that the expression level is simulated from $y_{gi} \overset{iid}{\sim} \text{UNIF}(M - A, M + A)$; and $1 - q$% chance that the expression level is simulated independently based on Equation 1 under $H_0 : A = 0$ (i.e., $y_{gi} = M + \varepsilon_{gi}$, $\varepsilon_{gi} \overset{iid}{\sim} N(0, \sigma^2)$). Figure S4b shows that the cosinor method achieves accurate nominal type I error rate control (i.e., 5%), showing robustness to outliers.

3. Correlated gene structure Instead of assuming all genes are independent, we simulate every 50 genes as a gene module with correlation coefficient $\rho$. The error term for each gene module $\boldsymbol{\varepsilon}_{50} \sim N(\mathbf{0}, \boldsymbol{\Sigma}_{50})$ where $\boldsymbol{\Sigma}_{50}$ is a symmetric matrix with diagonal elements being $\sigma^2$ and off-diagonal elements being $\sigma^2\rho$. We simulate a grid of $\rho = (0, 0.25, 0.5, 0.75)$. Figure S4c shows that when genes are correlated, the cosinor method maintains accurate nominal type I error rate control (i.e., 5%).

### S4 Non-parameteric curve fitting simulation setting

To explore the impact of number of time points per cycle NT on the goodness-of-fit for a sinusoidal wave, we simulate expression data from the sinusoidal model and perform non-parametric curve fitting. To be specific, we first choose the ZT points within a cycle to be $N_T = 2, 3, 4, 6, 8, 12$. Then for each $N_T$, we simulate $n = 48$ samples, and evenly allocated them at $2N_T$ time points across 2 full cycles (every $24/N_T$ hours, from -12h to 36h), resulting in $24/N_T$ samples at each time point. The expression values of samples at each time point $t_j$ are simulated independently from Equation 1, where we set $A = 1$, $M = 0$, and $\sigma = 1$. The LOESS regression is then used to fit a smooth curve through the data points. The LOESS regression is a nonparametric method using locally-weighted regression to fit a smooth curve over a scatter plot [1] . Such LOESS regression represents the smooth curve fitting without the sinusoidal assumption, which could reflect the minimum of $N_T$ that is necessary to capture the sinusoidal wave curve. The rationale for 2 full cycles is to improve the boundary behavior of the curve fitting within one cycle. In addition, to evaluate the effect of the phase shift, we set $\phi = 0, 1, \ldots, \min(24/N_T - 1, 6)$. By comparing the fitted non-parametric curves with the underlying sinusoidal wave, we can observe the minimum of NT that is necessary to capture the sinusoidal wave curve. The data points and fitted smooth curves in one circadian cycle [0, 24] are shown in Figure S2.

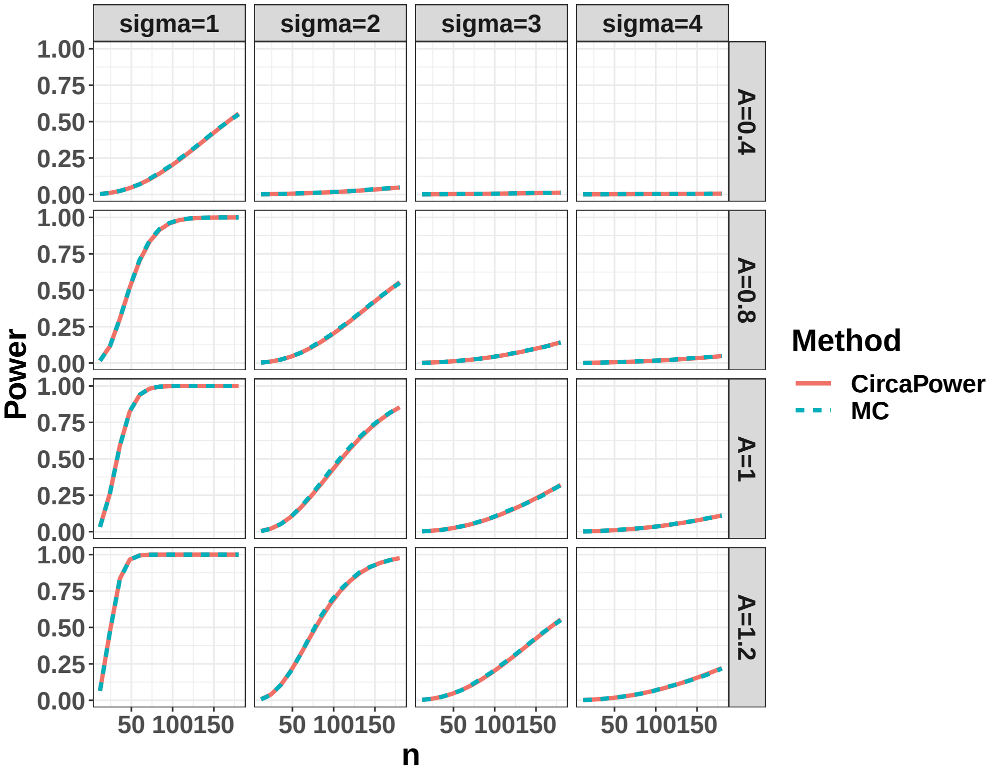


(a)

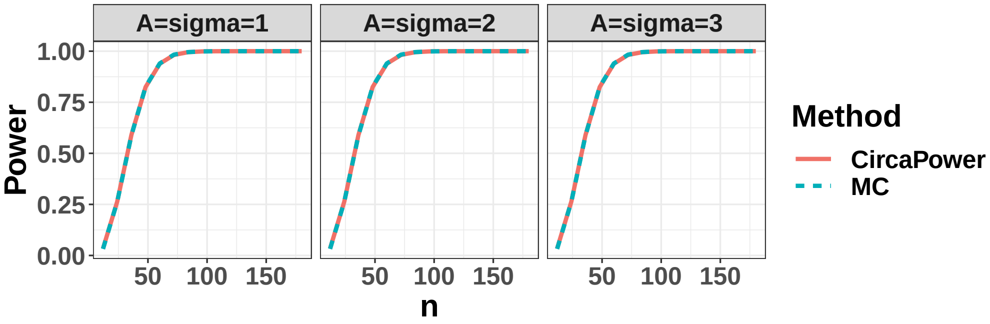


(b)

Figure S1: Compare power trajectories from the CircaPower with the Monte-Carlo (MC) algorithm. The red color curve denotes the CircaPower method, and the cyan color curve denotes the MC approach. (a) We vary the amplitude $A = 0.4, 0.8, 1, 1.2$ and $\sigma = 1, 2, 3, 4$. (b) We co-vary $A$ and $\sigma$ simultaneously (i.e., $A = 1, 2, 3$, $\sigma = 1, 2, 3$) while keeping their ratio as a constant (i.e., $r = A/\sigma = 1$).

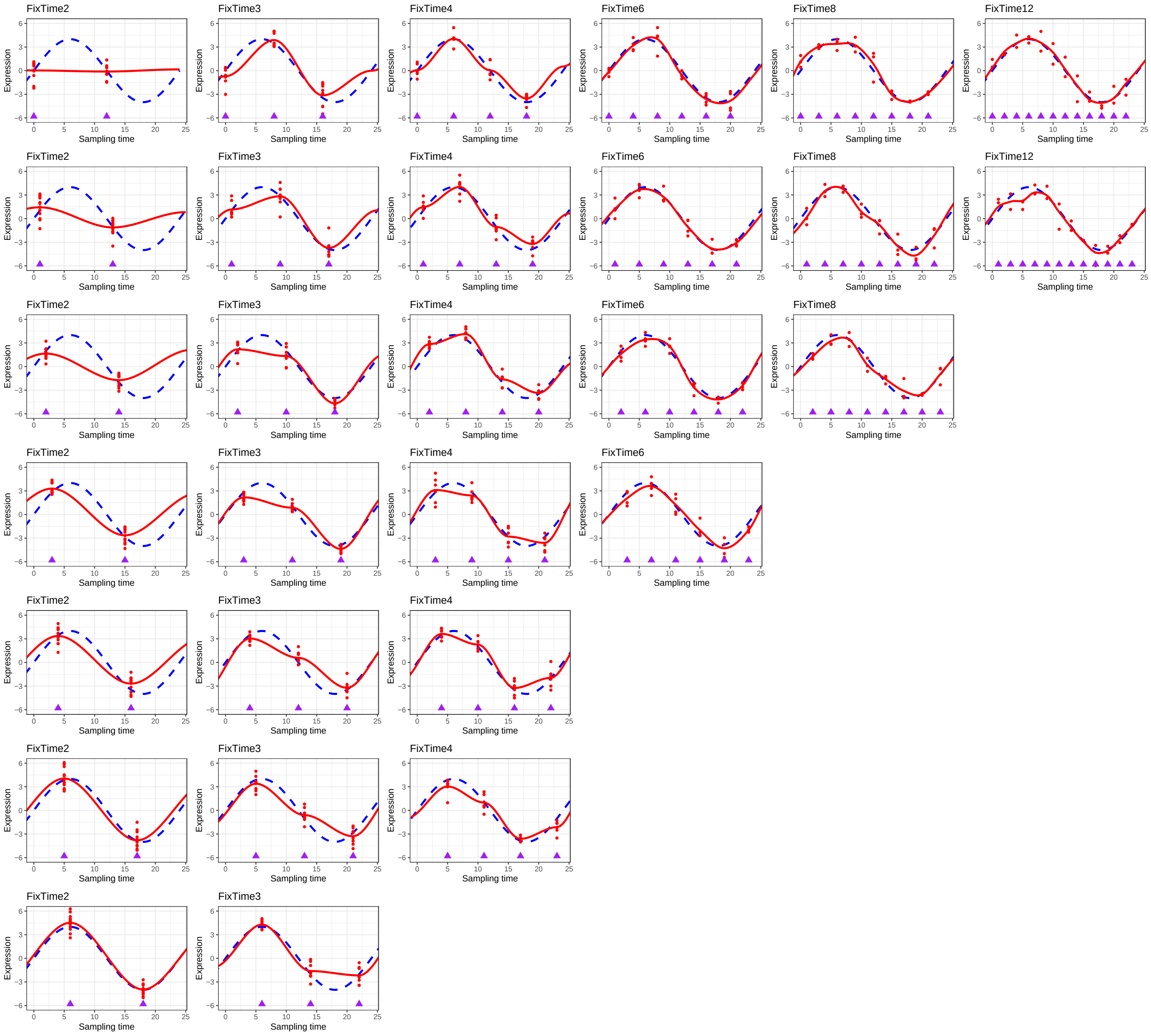


Figure S2: Non-parametric curve fitting to explore the impact of number of ZT points per cycle (i.e., $N_T$) on the goodness of fit for a sinusoidal wave. We choose the ZT points within a cycle to be $N_T = 2, 3, 4, 6, 8, 12$ (on columns). Then for each $N_T$, we simulate $n = 48$ samples evenly allocated at $2N_T$-time points across 2 full cycles (every $24/N_T$ hours, from -12h to 36h), resulting in $24/N_T$ samples at each time point. The rationale for 2 full cycles is to improve the boundary behaviour of the curve fitting within one cycle. The purple rectangles indicate sampling time points. The expression values of samples at each time point $t_j$ are simulated independently from Equation 1 with $A = 1$, $M = 0$, and $\sigma = 1$. The LOESS regression is used to fit a smooth curve through the all data points. To evaluate the effect of the phase shift, we set $\phi = 0, 1, \ldots, min(24/N_T - 1, 6)$ (on rows). The data points and fitted smooth curve in one circadian cycle [0, 24] are shown in this figure, which represents the smooth curve fitting without the sinusoidal assumption.

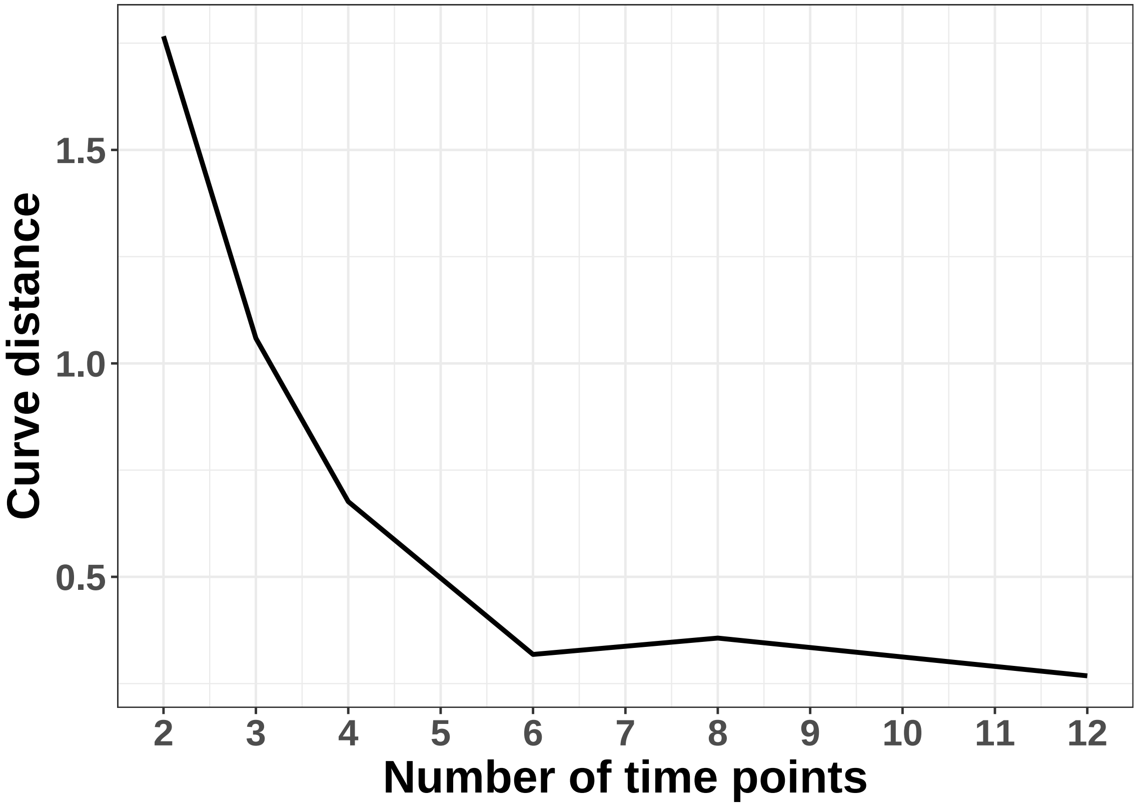


Figure S3: Elbow plot to evaluate the effect of number of ZT points per cycle (i.e., $N_T$) on the goodness of fit for a sinusoidal wave. The curve distance is calculated as $\sum_x \sqrt{\frac{(f(x)-\hat{f}(x)}{1000}}$ where $x$ are 1000 points evenly spaced between 0 and 24, $f(x)$ is the underlying sinusoidal curve evaluated at $x$ and $\hat{f}(x)$ is the fitted non-parametric curve evaluated at $x$. To avoid randomness, for each $N_T$, the non-parametric fitting is repeated 100 times and the mean curve distance is plot against the $N_T$.

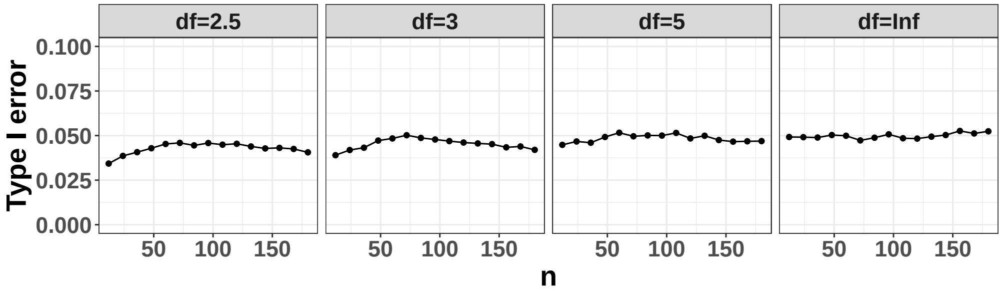


(a)

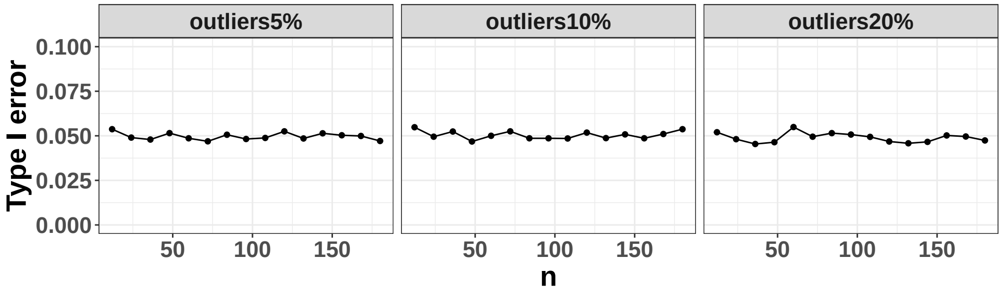


(b)

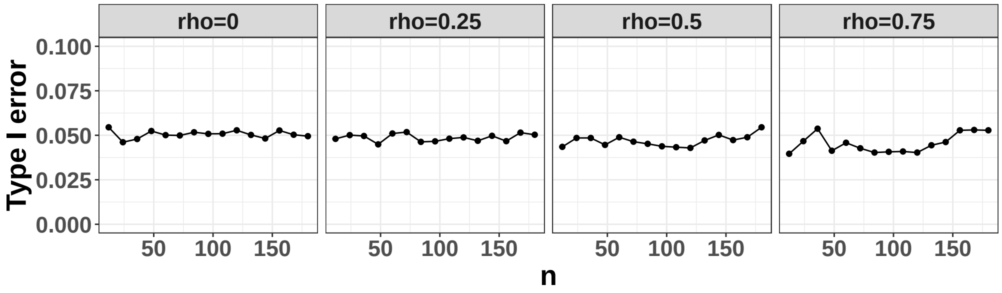


(c)

Figure S4: Actual type I error rate for the cosiner method in detecting circadian rhythmicity pattern at 5% nominal $\alpha$ level when there exist violation of model assumptions (i.e., independent Gaussian errors.) (a) shows the case when the error term comes from a heavy tail $t$ distribution (i.e., $\epsilon_i \sim t(df)$, $df$ is the degree of freedom of the $t$ distribution); (b) shows the case when there exists outliers; (c) shows the case when the errors of multiple genes are correlated (i.e., $(\epsilon_1, \ldots, \epsilon_m) \sim \text{MVN}(0, \Sigma)$, $m$ is number of correlated genes, and $\Sigma$ is the variance-covariance matrix for the multivariate normal distribution).

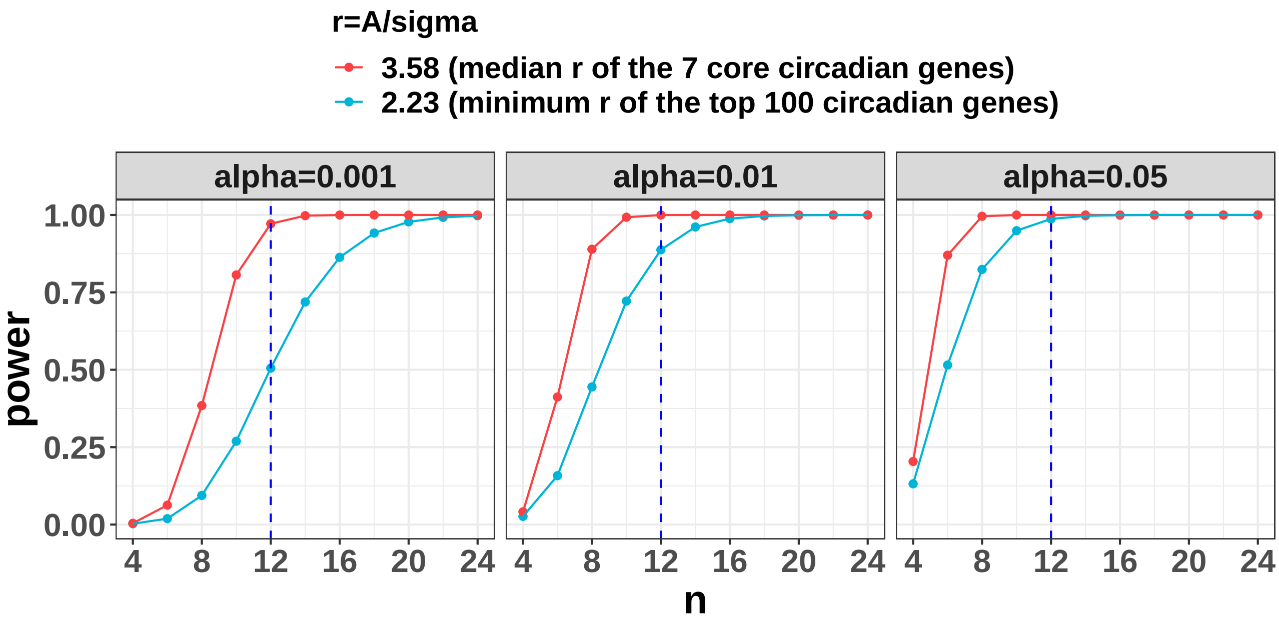


Figure S5: Case study: power trajectories using mouse muscle gene expression circadian data as the pilot data. The pilot data contains 24 mouse muscle samples collected every 2 hours across 2 full cycles. CircaPower is used to calculate the power. The intrinsic effect sizes were used as (i) median $r$ of the 7 core circadian genes; (ii) minimum $r$ of the top 100 significant circadian genes. By assuming different $\alpha$ to be 0.05, 0.01, 0.001, the power trajectories with respect to sample size is shown in this figure. The blue dashed lines indicate the detection power when $n = 12$ per cycle which is statistically equivalent to sampling every two hours across two full cycles.
